# *Streptococcus* Phage Genomes Reveal Extensive Diversity, New Taxonomic Insights, and Novel Endolysin-Derived Antimicrobial Peptides

**DOI:** 10.1101/2024.10.31.621281

**Authors:** Daniyal Karim, Shakhinur Islam Mondal, Mohimenul Haque Rolin, Nurnabi Azad Jewel, Hammadul Hoque, Colin Buttimer, Md Mahfuzur Rahman, Abul Kalam Azad, Arzuba Akter

## Abstract

The global rise of antibiotic-resistant bacteria, particularly among *Streptococcus* species, poses an escalating public health threat. Traditional antibiotic development has proven inadequate, making innovative approaches such as bacteriophage-based therapies promising alternatives. A deep understanding of phage biology at the genomic level is essential for advancing therapeutic applications. Here, we analyzed 709 *Streptococcus* phage genomes to bridge gaps in genomic diversity and propose revisions to *Streptococcus* phage taxonomy. The phage genomes were clustered based on shared proteins, resulting in 66 clusters and 35 singletons with significant variation in genome characteristics. Through proteome phylogeny, average nucleotide identity, and inter-cluster core genes, we propose 21 new family-level classifications and 296 genus-level subclusters, providing an updated framework for *Streptococcus* phage taxonomy. Further analysis revealed diverse domain architectures in *Streptococcus* phage endolysins, including previously unreported structures. Specific domains were associated with distinct streptococcal hosts, suggesting adaptive evolution. We also observed variation in endolysin gene organization, with purifying selection acting on most sites, though some were subject to diversifying selection. Additionally, 182 novel endolysin-derived antimicrobial peptides (AMPs) were identified, some exhibiting antifungal, antiviral, cell-penetrating and non-toxic properties. Molecular dynamics and docking simulations demonstrated high stability and strong binding affinity of peptides EP-39 and EP-121 to the *Streptococcus pneumoniae* virulence factor autolysin. This is the first comprehensive comparative study of *Streptococcus* phage genomes, providing critical insights into phage diversity and taxonomy. It also highlights the therapeutic potential of endolysin-derived AMPs against multidrug-resistant *Streptococcus* strains. Further experimental validation is required to assess their clinical potential.

## 1. Introduction

The widespread emergence of antibiotic-resistant bacteria, driven by the overuse and misuse of antibiotics in both medicine and agriculture, poses a critical global health threat (1). In the United States alone, antimicrobial resistance (AMR) accounts for 99,000 deaths annually from hospital-acquired infections, with global projections estimating 10 million deaths by 2050 if current trends continue (2). Among the key pathogens contributing to this crisis are various *Streptococcus* species, including Group A *Streptococcus* (GAS) such as *S. pyogenes*, and Group B *Streptococcus* (GBS) such as *S. agalactiae*, which are significant contributors (3,4). GAS leads to severe conditions like necrotizing fasciitis and rheumatic fever (5), while GBS is a major cause of neonatal infections, including meningitis and sepsis (4). Additionally, *Streptococcus mutans* and *Streptococcus pneumoniae* are implicated in serious human infections (6,7). The rapid spread of resistance genes, particularly in *S. pneumoniae*, exacerbates treatment challenges (8,9).

Given the limitations of traditional antibiotic development, there is a growing need for unconventional approaches to combat antibiotic-resistant bacteria (10,11). Among the alternative strategies being explored, bacteriophage-based therapies have garnered significant attention (12,13). Bacteriophages, viruses that infect bacteria, utilize lysogenic or lytic cycles. The lytic cycle, mediated by phage-encoded peptidoglycan hydrolases (PGHs) named endolysins, degrades the bacterial cell wall, causing rapid bacterial cell death (13–15). Phage-based therapy leverages this property by administering phages or their derived products to target and kill specific bacteria. Endolysins are particularly promising as therapeutic agents due to their specificity, low resistance risk, and synergy with other antibacterial agents (16,17). In addition to endolysins, bacteriophage PGH-derived antimicrobial peptides (AMPs) have also gained interest in combating antibiotic-resistant pathogens (18–20). Antimicrobial peptides (AMPs) are low molecular-weight oligopeptides, 15–150 amino acids in length, that exhibit broad-spectrum antimicrobial activity by disrupting bacterial cell membranes (21–23). Bacteriophage-based therapies have been tested against various *Streptococcus spp.* For instance, the Eliava Institute effectively used phages to treat bacterial pathogens, including *Streptococcus spp*., with 74% of bacterial dysentery patients showing improvement (24). Endolysins, such as PlyC, PlyPy, and pneumococcal variants like Pal and Cpl-1 have shown great promise against streptococcal infections. PlyC effectively lyses groups A, C, and E *streptococci*, while PlyPy, derived from *S. pyogenes*, targets a broader range of species, including *S. uberis* and *S. gordonii* (25,26). Pneumococcal endolysins like Pal and Cpl-1 have demonstrated efficacy in treating pneumonia and bacteremia in vivo (27–29), and engineered endolysins such as Cpl-7s, Cpl-711 and ClyJ have also shown promise in vivo (30–32). While PGH-derived AMPs have not yet been tested against *Streptococcus* species, they have demonstrated efficacy against other pathogens like *Acinetobacter baumannii* and *Pseudomonas aeruginosa* (18–20,33).

The growing accessibility of completely sequenced phage genomes has facilitated *in silico* investigations that provide a comprehensive understanding of phage biology, evolutionary trends, and host-phage interactions. These studies are crucial for identifying therapeutically valuable phages and phage-derived proteins. Comparative genomic analyses have been conducted on phages that infect *Mycobacterium*, *Acinetobacter*, *Staphylococcus*, *Pseudomonas*, *Salmonella*, *Vibrio cholerae*, *C. difficile* and Alphaproteobacteria (34–41). For *Streptococcus* phages, such analyses have primarily focused on dairy phages, particularly those targeting *Streptococcus thermophilus*, as well as on ten temperate *Streptococcus pneumoniae* phages (42–44). Previous *in silico* studies on phage-derived endolysins have explored domain diversity within available dsDNA bacteriophage endolysins in databases (45). A similar analysis was conducted on protein domain sequences in phage and prophage endolysins (46), and a subsequent study identified putative endolysins from uncultured bacteriophages, revealing new domains (47). More recent studies have examined the domain diversity, host association, and molecular evolution of the lytic arsenal of *Pseudomonas* and *Stenotrophomonas* phages (48,49). Our recent study on *Streptococcus pneumoniae* phage endolysins identified previously unreported additional cell wall-binding repeats (50). Additionally, a database called PhaLP was developed to investigate phage endolysins and their evolution (51). *In silico* studies on phage PGH-derived AMPs have led to the development of databases such as ESKtides, which specifically mines and catalogues AMPs derived from ESKAPE phages (52). However, no comparative studies have yet been conducted on all sequenced *Streptococcus* phage genomes, the domain diversity and evolution of their endolysins or the mining of AMPs from these endolysins.

This study addresses these gaps by conducting an extensive analysis of all sequenced *Streptococcus* phage genomes using a variety of computational approaches. The genomes were grouped into clusters and subclusters, establishing genus and family-level relationships among the phages. Additionally, we examined the prevalence and diversity of domain architectures within *Streptococcus* phage endolysins and their evolutionary trajectories. Finally, we mined AMPs from these endolysins and explored their interactions with the virulence factor autolysin of *Streptococcus pneumoniae*.

## 2. Methods

### 2.1 Retrieval of phage genome sequence and annotation

*Streptococcus* phage genome sequences were downloaded from the INPHARED database (July 21, 2023) (https://github.com/RyanCook94/inphared) (53). To remove duplicate phage genome sequence, CD-HIT-EST (sequence identity threshold = 1, word size = 10) was employed (54). To predict the coding sequencing (CDS) of those non-redundant phage genomes, Prodigal implemented in Pharokka version 1.3.2 was used in meta mode. PHROGs database is utilized for the functional annotation of the predicted CDS. Pharokka additionally employs tRNAscan-SE 2 for tRNA prediction, Aragorn for tRNA prediction and MinCED for CRISPR prediction. The Comprehensive Antibiotic Resistance Database (CARD) and the Virulence Factor Database (VFDB) are concurrently utilized to identify antimicrobial resistance and virulence genes in Pharokka (55).

### 2.2 Genome Clustering, pangenome analysis and Taxonomy

Using PhaMMseqs (35% amino acid identity and 80% coverage), proteins of all genomes were assigned to phage protein families or phams with the pangenome option (-p) enabled (56). *Streptococcus* phages were clustered using PhamClust (clustering threshold = 0.3) based on shared phams (57). To calculate the intergenomic similarity of phage genomes, VIRIDIC was used in default parameters. To assign cluster members into subclusters, a 70% average nucleotide identity cutoff value was used (58). To identify core genes within cluster members and intercluster members, Pirate (30% identity, 50% coverage) was used (59). ViPTreeGen was used to generate a proteomic tree of viral sequences based on genome-wide sequence similarities along with all current members of *Caudoviricetes* except *Crassvirales* as reference sequence (n = 3,823, after duplicate removal, April 25, 2023) (60). The proteomic tree was visualized and annotated on iTOL v6 server (61).

### 2.3 Endolysin prediction, Phylogeny and Selection Pressure analysis

To obtain putative endolysin, PhaMMseqs assigned phams that contained endolysins were identified using “endolysin” as the keyword. Then, putative endolysin sequences were annotated for domain information using HMMER v3.4, with Pfam-A.hmm as the database (http://hmmer.org/). The putative endolysin sequences were grouped based on their domain architecture as well. The function “build” of ETE3 3.1.2 as implemented on the GenomeNet was used to align and reconstruct the phylogeny of the curated endolysins (https://www.genome.jp/tools/ete/) (62). MAFFT v6.861b was utilized to conduct alignment with the default settings (63). The alignment that was obtained was cleaned up with the gappyout algorithm in trimAl v1.4.rev6 (64). The ML tree was inferred using IQ-TREE 1.5.5 ran with ModelFinder and tree reconstruction. The best-fit model according to BIC was WAG+F+R5. Tree branches were tested by SH-like aLRT with 1000 replicates (65). The phylogenetic tree was visualized through the iTOL v6 server (61).

Before performing selection pressure analysis on each endolysin pham, containing at least 10 unique endolysin sequences, stop codons were removed and nucleotide sequences were aligned using MACSE v2 (66). Using Datamonkey webserver, recombination and selection pressure analysis was performed utilizing the following programs-GARD was used to identify recombination breakpoints and for accurate selection inference before usage of MEME, FUBAR, SLAC, FEL in default parameters (67–70).

### 2.5 Antimicrobial peptide identification and characterization

To fragment each endolysin into peptides, a sliding window-based approach was applied iteratively over the full length of its amino acid sequence. A window size of 20 amino acids was chosen and the window moved one amino acid at a time. Six AMP prediction tools were used to screen these peptides for their antimicrobial properties. These were AMPlify, amPEPpy 1.0, AxPEP, AmpGram, CAMPR4, and Antimicrobial Peptide Scanner vr.2 (71–76). Peptides which were predicted to have antimicrobial properties by all six tools were selected for further analysis. dPABBs were utilized to predict the antibiofilm capabilities of these putative AMPs, whilst Antifp and iAMPred were utilized to predict their antifungal properties (77–79). iACVP and iAMPred were utilized to predict antiviral activity (79,80). ToxIBTL was used to determine the toxicity of the selected AMPs (81). BChemRF-CPPred, MLCPP2.0 were utilized to examine the cell-penetrating characteristics of those putative AMPs (82,83).

### 2.6 Structural Modelling and Peptide stability assessment

AlphaFold2 was used to generate the 3D structure of the putative AMPs (84). To check the structural stability of the AMP, molecular dynamic simulation was employed by GROMACS (2023.3 release) using CHARMM36 all-atom force field (85,86). After solvating each AMP with the CHARMM-modified TIP3P water model, an appropriate amount of Na/Cl ions was added to neutralize the solution. The system’s energy was minimized through the implementation of a steepest descent minimization algorithm consisting of 50,000 steps, followed by a 2-phase equilibrium step. The first equilibration phase was performed using an ensemble consisting of a constant number of particles, volume, and temperature (NVT). In contrast, the second equilibration phase was executed using an ensemble consisting of a constant number of particles, pressure, and temperature (NPT) for a duration of 50,000 steps (100 ps) through the utilization of the Particle Mesh Ewald (PME) method. Finally, molecular dynamics simulations of each AMPs were carried out for 50 nanoseconds (ns). Root mean square deviation (RMSD) values were calculated from the simulation trajectory file using gmx rms in Gromacs.

### 2.7 Molecular docking

Based on the lowest average RMSD value, the top thirty AMPs were selected for molecular docking analysis. To explore the binding interactions between newly predicted AMPs with virulence factors of *Streptococcus pneumoniae,* one protein target was chosen namely autolysin (PDB ID: 4X36). To predict the binding sites of this protein target, the CASTp server was used (87). The molecular docking of the top 30 AMPs with the selected protein target was performed using HADDOCK server (88).

### 2.8 Molecular dynamics simulation

Molecular dynamics (MD) simulation analysis was performed for 100 nanoseconds on the top three autolysin-AMP complexes, as determined by the HADDOCK score. GROMACS built-in programs were utilized to examine RMSD (Root Mean Square Deviation), RMSF (Root Mean Square Fluctuation), and Rg (Radius of gyration), SASA (solvent-accessible surface area) of autolysin and peptide using molecular dynamics simulation trajectories. The graphs of the analyses were generated by xmgrace. Gmx_MMPBSA tool was used to determine end-state free energy over the last 30 ns of MD simulation of autolysin-AMP complexes (89).

## 3. Result

### 3.1 Diversity of *Streptococcus* phages

As of July 21, 2023, the INPHARED database contained 922 *Streptococcus* phage genomes. After removing duplicates, 709 genomes were selected for this study. These phages belong to the class *Caudoviricetes*. The majority of the phages (n = 580) remain unassigned to a family, while 126 belong to *Aliceevansviridae*, one to *Rountreeviridae*, and two to *Salasmaviridae*. At the genus level, the phages are classified as follows: *Brussowvirus* (n = 34), *Vansinderenvirus* (n = 11), *Fischettivirus* (n = 1), *Saphexavirus* (n = 1), *Moineauvirus* (n = 81), *Paclarkvirus* (n = 12), *Piorkowskivirus* (n = 7), *Stonewallvirus* (n = 1), and *Cepunavirus* (n = 2), with 559 phages remaining unassigned.

Genome lengths range from 11 kb to 110 kb (Figure 1). Phages from Rountreeviridae and Salasmaviridae have smaller genomes, while those from Aliceevansviridae average around 36 kb. GC content varies between 33% and 45%, with no significant correlation to genome length (Pearson R = 0.0029). Genome length moderately correlates with coding sequences (CDS) (Pearson R = 0.7286) but had negligible correlation with coding density (Pearson R = 0.0033) (Figure S1). ARAGORN identified 93 transfer RNAs (tRNAs) in 78 genomes: 64 had one tRNA, 13 had two, and one had three.

**Figure 1:**
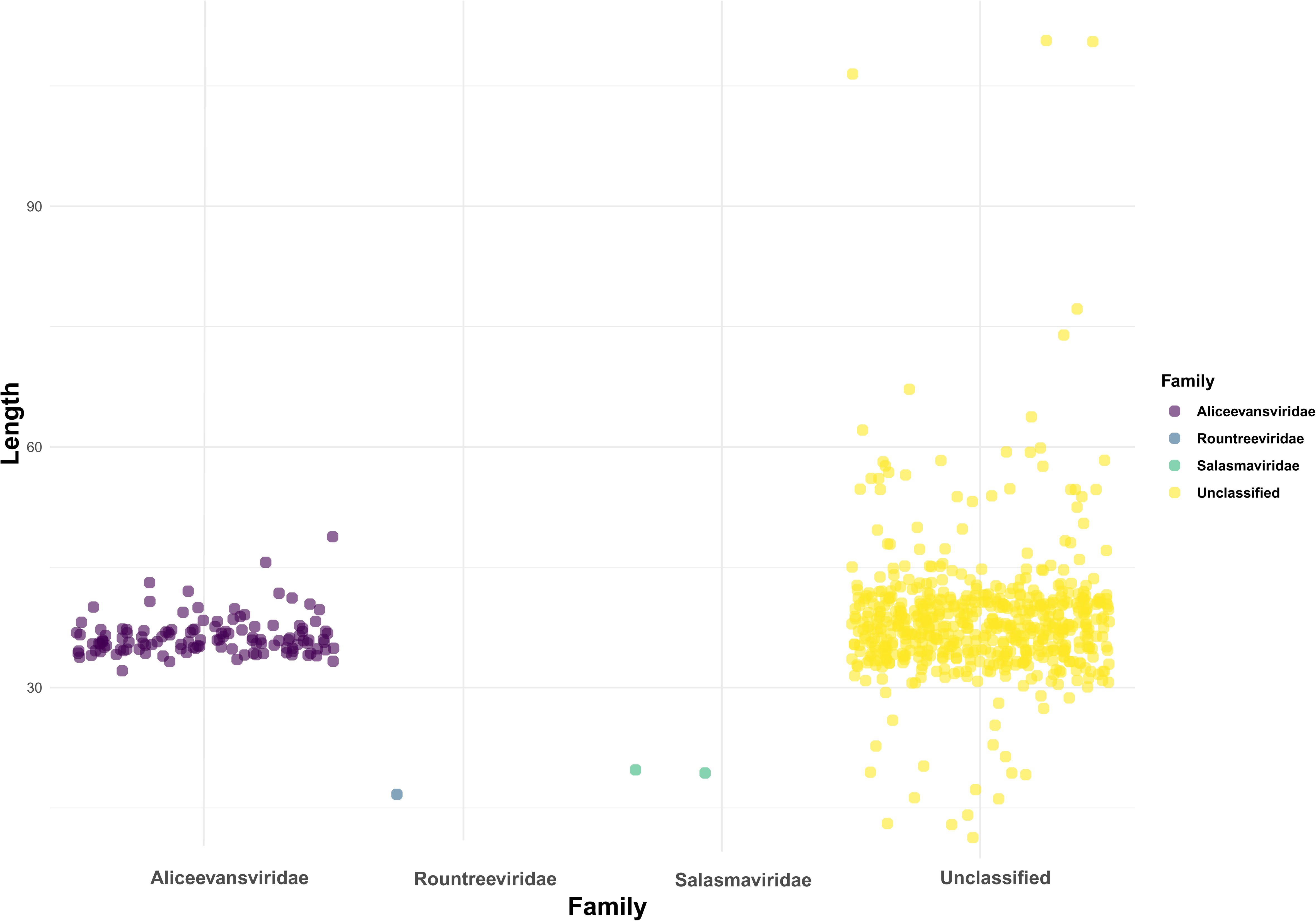
Jitter plot of genome length of *Streptococcus* phages. In X-axis, *Streptococcus* phages belonging to ICTV classified families are shown and in Y-axis, genome length is shown. High genome length variation is observed in unclassified phages.

CARD detected 62 antimicrobial resistance genes (ARGs) in 24 genomes, with macrolide-resistance gene *mel* (n = 13) being the most common, followed by *ANT*(*6*)*-Ia* (n = 6) and *AAC6_Ie_APH2_Ia* (n = 5), indicating aminoglycoside resistance.

Notably, phage phi-SsuZKB4_rum had the most resistance genes (n = 7), followed by phi-SsuSC05017_rum (n = 5), phi-SsUD.1 (n = 5), and phiSC070807 (n = 5). The largest genomes, including phi-SgaBSJ31_rum, phi-SgaBSJ27_rum, and phi-SsuZKB4_rum, also carried resistance genes. VFDB revealed 279 virulence factor genes in 186 genomes. Many of the *Streptococcus* phages with virulence factor genes were isolated from *Streptococcus pneumoniae* (n = 73) or *Streptococcus pyogenes* (n = 66). Among these 186 genomes, 69 contained multiple virulence factor genes. The most prevalent virulence factor gene was autolysin, found 84 times, followed by hyaluronidase with 69 occurrences. Additionally, various streptococcal exotoxin precursor genes, including *SpeA, SpeC, SpeG, SpeH, SpeI, Spek, SpeL,* and *SpeM*, were observed with a total of 54 occurrences. MinCED identified two CRISPR spacer sequences within the genomes of phages 128 and 142. Detailed information on the general characteristics of these phages (Table S1), as well as the antimicrobial resistance genes identified by CARD (Table S2) and the virulence factors identified by VFDB (Table S3), can be found in the supplementary table.

### 3.2 Cluster Assignment and Taxonomy of *Streptococcus* Phages

To analyze the taxonomy of 709 *Streptococcus* phage genomes, we used protein sharing as the primary clustering criterion. PhamClust calculated the Proteomic Equivalence Quotient (PEQ) between phage pairs based on shared gene identity, with PEQ values ranging from 0% (no shared genes) to 100% (complete identity) (Table S4). This approach identified 66 distinct clusters (Cluster 1–66), along with 35 singletons (Figure 2). Additionally, a 70% Average Nucleotide Identity (ANI) threshold yielded 296 subclusters within these clusters (e.g., Cluster 1A, 1B) (Table S5).

**Figure 2:**
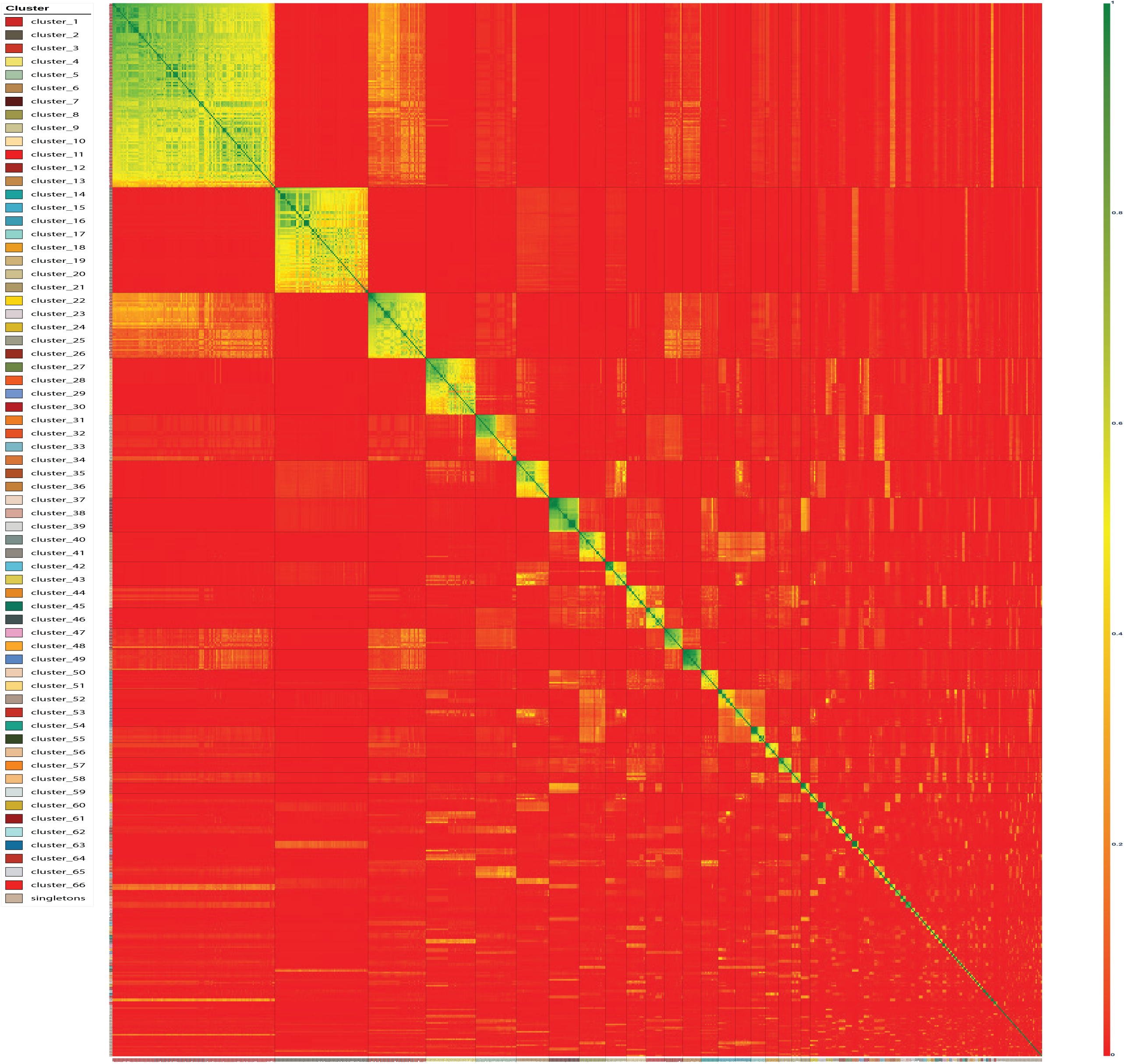
Heatmap of PEQ values among the *Streptococcus* phage genomes. Based on PEQ values, 709 *Streptococcus* phage genomes were assigned to 66 clusters and 35 singletons. Phage genome accessions are colored based on cluster identity and clusters which had at least 10 phages (Cluster 1 – Cluster 19) are marked using horizontal and vertical lines.

Cluster 1 was the largest with 124 members, while clusters 20 to 66 contained fewer than 10 members each. Cluster 2, containing 71 phages, included three large phages (phi-SgaBSJ31_rum, phi-SgaBSJ27_rum, phi-SsuZKB4_rum) with genomes ranging from 33 kb to 110 kb and encoding between 37 to 112 genes. Although phi-SgaBSJ31_rum and phi-SgaBSJ27_rum shared high genomic similarity (Figure 3A), phi-SsuZKB4_rum (∼110 kb genome length) shared higher genomic similarity with phi-SsuSC05017_rum (∼57.5 kb genome length) and phiSC070807 (∼57.6 kb genome length), despite significant genome length differences (Figure 3B). This is also verified by the proteomic tree discussed later (Figure S2). Cluster 18 exhibited similar variability, with Javan158 (∼19.4 kb) encoding 25 genes and Javan141 (∼42.4 kb) and Javan149 (∼41.7 kb) each encoding 70 genes. Genome comparison and PEQ values indicated that Javan158 shared genomic similarity with Javan141 (Figure S3).

**Figure 3:**
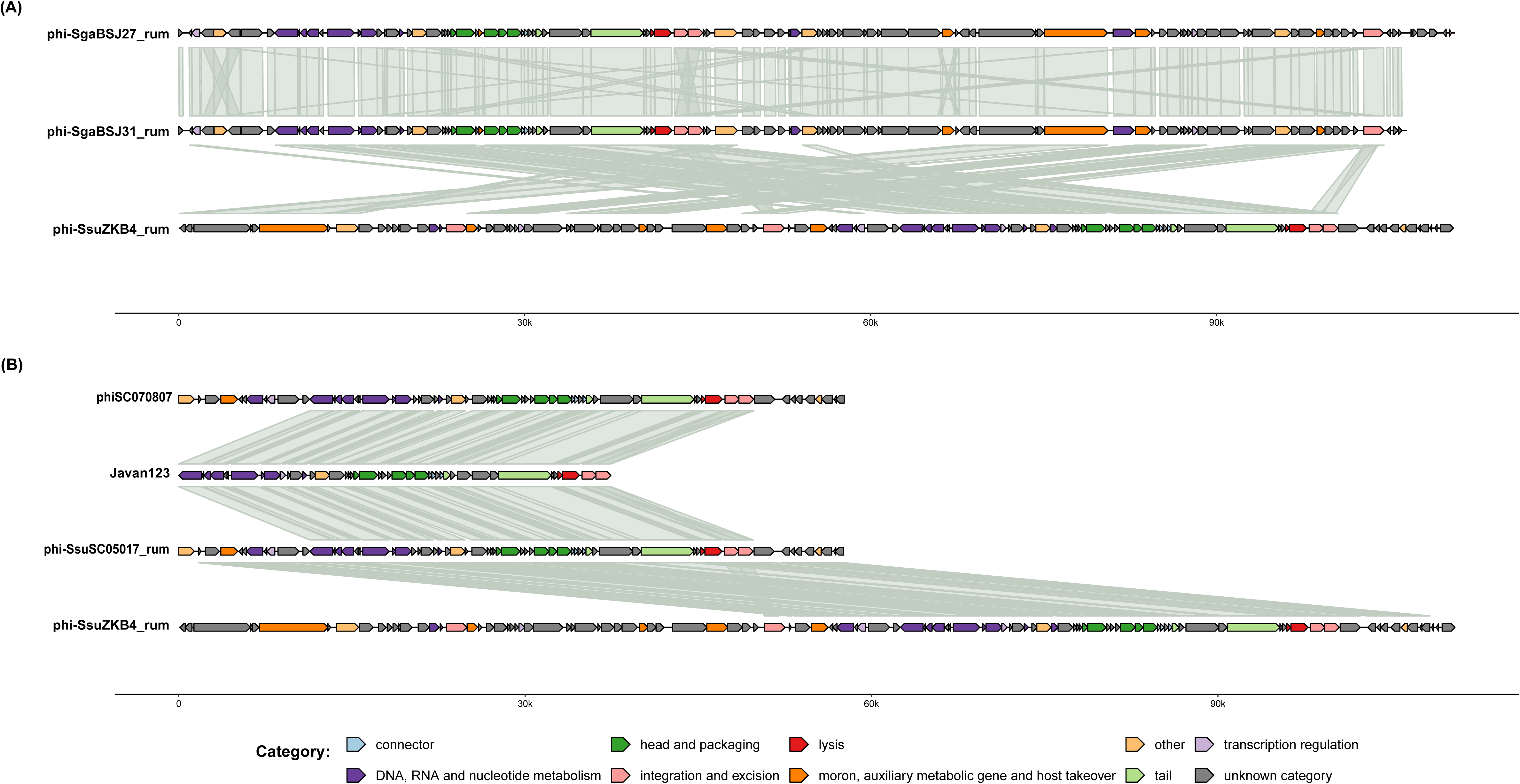
Genome comparison among all three large *Streptococcus* phages: phi-SgaBSJ31_rum, phi-SgaBSJ27_rum, phi-SsuZKB4_rum and genome comparison among phi-SsuZKB4_rum, phi-SsuSC05017_rum, Javan123 and phiSC070807. A) Genome comparison among all three large *Streptococcus* phages showed that phi-SgaBSJ31_rum and phi-SgaBSJ27_rum is highly conserved and similar. B) In contrast, the other large phage, phi-SsuZKB4_rum, showed high conservation with phi-SsuSC05017_rum. The genes are colored according to their functional categories as indicated in the legend.

A total of 37,567 proteins from these 709 genomes were grouped into 3,455 phams (protein families) using PhaMMseqs, of which 1,480 (42.83%) were unique orphams (proteins found in only one genome). These orphams were found in 372 genomes, with 30 phages containing more than 10 orphams. The phage SP-QS1, a singleton, had the highest number of orphams (n = 88), followed by cluster 2’s phi-SsuFJNP8_rum (n = 25), Javan630 (n = 22), and phi-SsuFJZZ32_rum (n = 22). Singleton phages had the highest orpham count (n = 350), followed by phages in clusters 2 (n = 210), 5 (n = 61), and 1 (n = 60) (Figure S4). At a 30% amino acid identity threshold, PIRATE and PhaMMseqs found no shared core genes among all 709 phages.

A proteomic approach was used to determine family-level rank, along with an inter-cluster core gene-sharing approach. A hierarchically clustered tree (Figure S2) was generated based on pairwise tBLASTx scores using VipTreeGen to process the 709 *Streptococcus* genomes along with all current members of *Caudoviricetes* except *Crassvirales* (n = 3,823, after duplicate removal as of April 25, 2023). Clusters 1 to 66 formed distinct monophyletic clades within the proteomic tree, with exceptions in clusters 21, 41, and 64. PIRATE was utilized at a 30% amino acid identity threshold to examine the occurrence of core genes within and between clusters (Table S6).

Assessment of proteomic tree and inter-cluster core genes indicated inconsistencies in the ICTV taxonomy of *Streptococcus* phages. For example, *Aliceevansviridae* did not form a single clade but appeared across clusters 1, 3, and 12 (Figure S2). Cluster 1 members formed a monophyletic clade with clusters 65, 35, and 38, sharing 5 core genes among 135 genomes (Figure 4). Cluster 3 exhibited relatedness to clusters 42, 13, 14, and 31, sharing 4 core genes among 80 members (Figure S5). Cluster 12 formed a monophyletic clade with several other clusters, sharing 7 core genes among 94 members (Figure S6).

**Figure 4:**
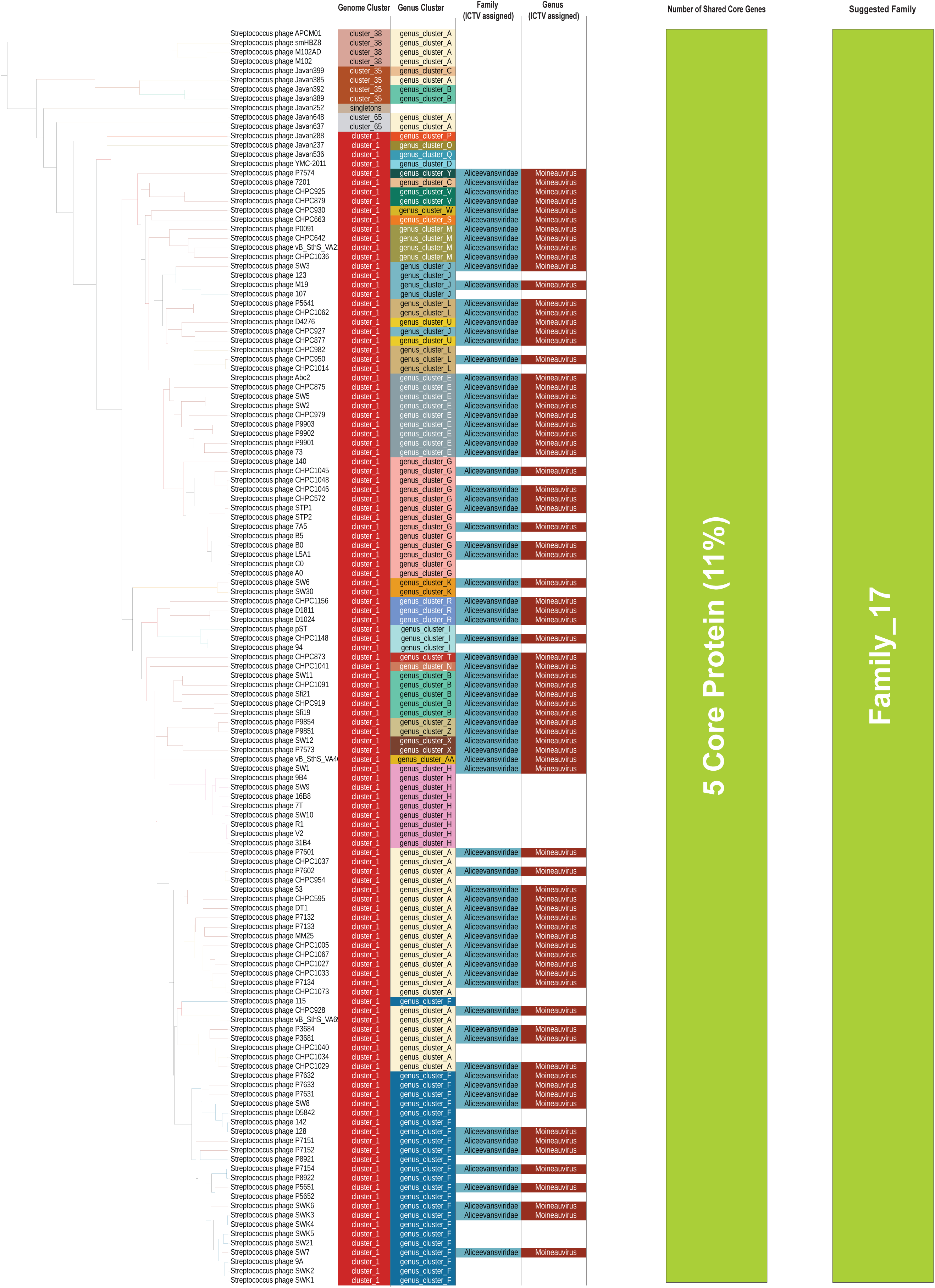
Fragment of the phylogenetic tree generated by ViPTreeGen using 709 *Streptococcus* phage sequences along with all current members of *Caudoviricetes* except *Crassvirales* (n = 3,823, after duplicate removal, April 25, 2023) showing cluster 1 formed monophyletic clade with cluster 65, 35, and 38 and shared 5 core genes among 135 genomes.

Based on the proteomic tree and shared core genes (at least four shared core genes), 21 new family-level ranks can be suggested, along with two previous families *Salasmaviridae* and *Rountreeviridae*. No family level rank could be suggested for phage SP-QS1, phiZJ20091101-2, phiJH1301-3 (Figure S2).

At the genus level, employing a 70% ANI cutoff within each cluster delineated 296 subclusters, each representing a unique genus. Notably, the ICTV’s classification of *Moineauvirus* and *Brussowvirus* phages was inconsistent, with *Brussowvirus* phages distributed across subclusters 3A to 3K (Figure S5), and *Moineauvirus* phages spanning subclusters 1A to 1AA, with some exceptions (e.g., 1D, 1P, 1O, and 1Q) (Figure 4). Additionally, some genus clusters did not form monophyletic clades; for instance, subcluster 1F phage 115 appeared within subcluster 1A, and subcluster 1J’s CHPC927 appeared between 1U members. Manual review showed that subcluster 1J phages had at least 70% ANI similarity with subcluster 1U’s CHPC877, but phages SW3 and M19 had lower ANI similarities (68.83% and 66.48%) with subcluster 1U’s D4276. Additionally, *Piorkowskivirus* phages appeared in subclusters 13A,13B and 13C (Figure S5). ANI values showed marginal differences between 13A and 13B subcluster members, with phage 9874 from 13B showing 69.9% and 69.3% similarity to phages SW 22 and SW 25 from 13A and phage CHPC926 from 13B showed at least 70% similarity with 13A subcluster members. On the contrary, phage CHPC577 from 13C showed less than 70% ANI similarity with most of the members of 13A and 13B. Proposed family and genus-level ranks, along with genome lengths, GC content, coding capacity, and shared core genes, are detailed in supplementary tables (Tables S7 and S8).

### 3.3 *Streptococcus* Phage Endolysin diversity and domain architectures

A total of 814 endolysins were identified in 694 *Streptococcus* phages, while 15 phages lacked endolysins. These 814 endolysins were clustered into 53 phams. Pham 63 was the largest (n = 109), followed by pham 72 (n = 104), pham 80 (n = 98), and pham 87 (n = 89); 38 phams contained fewer than 10 endolysins (Figure S7). Based on domain architectures and domain hits, 809 endolysins were classified into 54 groups, with five lacking significant domain hits (Figure 5). The largest group, group 1 (n = 195), featured Amidase_5 as the enzymatically active domain (EAD) and ZoocinA_TRD as the cell wall binding domain (CWBD). Group 2 (n = 89) contained endolysins with only ZoocinA_TRD as the sole CWBD. *Streptococcus* phage endolysins exhibited both mono-catalytic and bi-catalytic domain organizations (Table S9).

**Figure 5:**
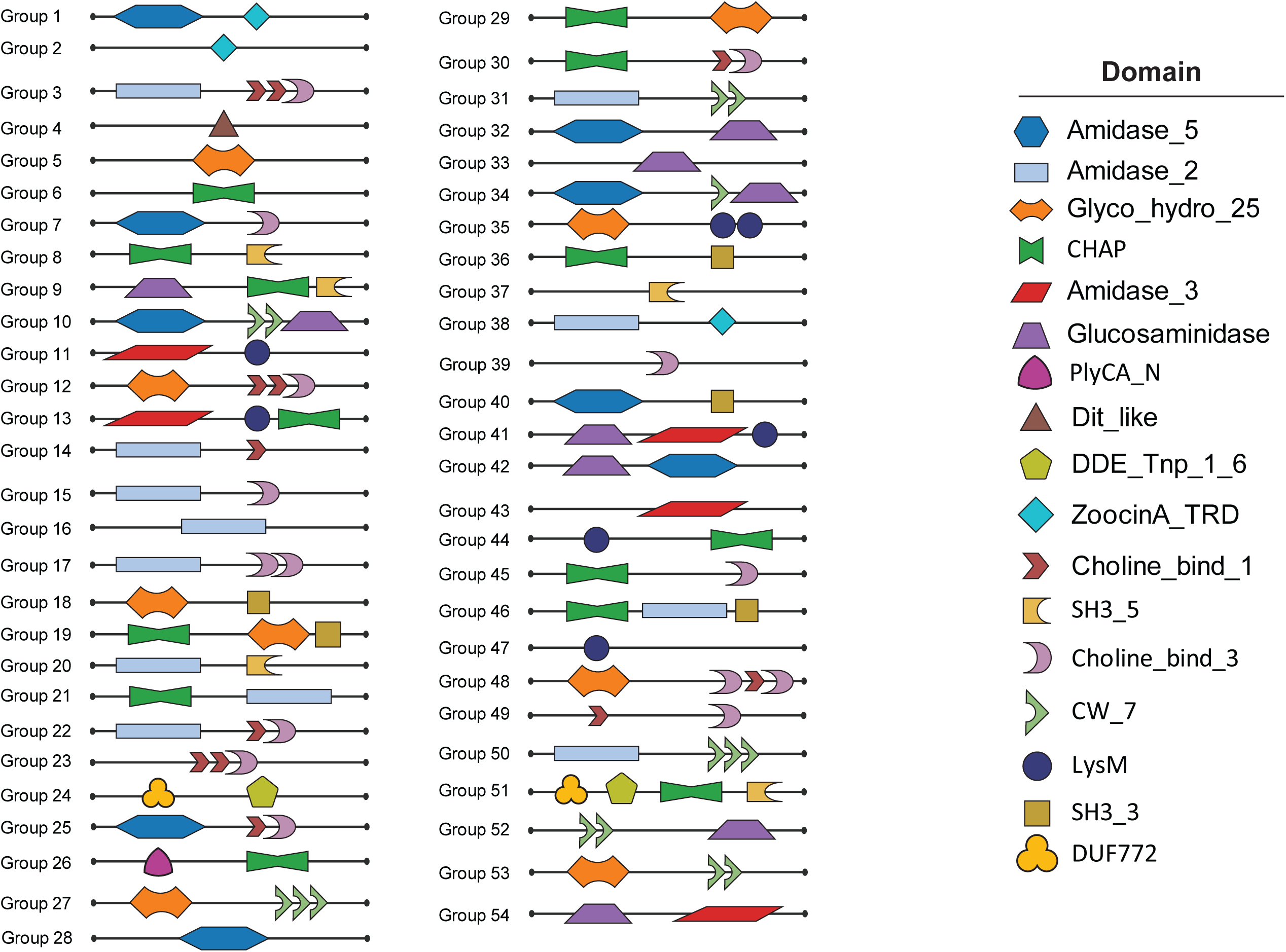
Observed domain architecture in 54 endolysin groups based on observed domain architectures and domain hits. Each shape indicates a single domain hit.

Domain architecture analysis against bacterial hosts revealed that mono-catalytic domain architecture with Amidase_5 as the EAD and ZoocinA_TRD as the CWBD was most common in *Streptococcus thermophilus* phage endolysins (n = 178). *Streptococcus pneumoniae* phages predominantly featured a similar mono-catalytic structure, with Amidase_2 combined with either two Choline_Bind_1 and one Choline_Bind_3 repeat or one Choline_Bind_3 and one Choline_Bind_1 repeat (n = 61 and n = 5, respectively). Another prevalent mono-catalytic form, featuring CHAP and SH3_5 domains, appeared in *Streptococcus pyogenes* (n = 10), *Streptococcus dysgalactiae* (n = 15), and *Streptococcus suis* (n = 19) phage endolysins. In bi-catalytic architectures, *Streptococcus suis* phages exhibited architectures like Amidase_3-LysM-CHAP (n = 24) and Amidase_5-CW_7-CW_7-Glucosaminidase (n = 14). *Streptococcus pyogenes* phages commonly displayed Glucosaminidase-CHAP-SH3_5 (n = 36) and Amidase_5-CW_7-Glucosaminidase (n = 10). CHAP-SH3_5 architecture was the most widespread, targeting 19 of 47 streptococcal hosts, followed by Amidase_3-LysM-CHAP (n = 12), Amidase_5-ZoocinA_TRD (n = 11), CHAP (n = 9), and Amidase_5-CW_7-CW_7-Glucosaminidase (n = 9) (Figure 6).

**Figure 6:**
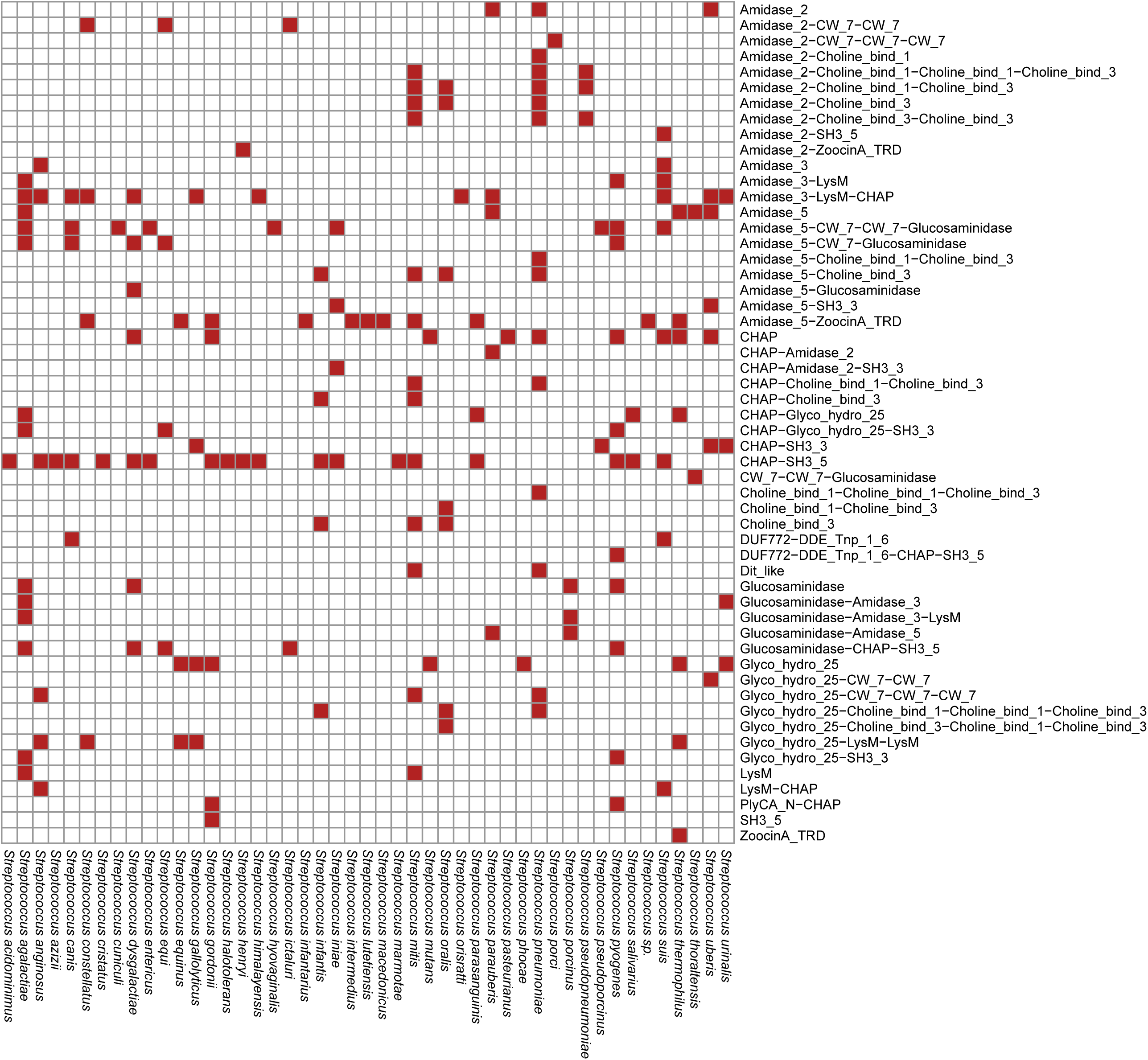
A heatmap illustrating the presence or absence of domain architecture against the 47 *Streptococcal* bacterial host. In X-axis, domain architectures are shown. Y-Axis indicates bacterial hosts. Red filled box represents presence of domain architecture.

When analyzing EADs against bacterial hosts, Amidase_5 was the most prevalent in *Streptococcus thermophilus* phage endolysins (n = 179). Amidase_2 dominated in *Streptococcus pneumoniae* (n = 76), while Amidase_3 appeared frequently in *Streptococcus suis* (n = 27) and *Streptococcus agalactiae* (n = 12). CHAP was abundant in *Streptococcus suis* (n = 47), *Streptococcus dysgalactiae* (n = 31), and *Streptococcus pyogenes* (n = 60), while Glucosaminidase was common in *Streptococcus pyogenes* (n = 53), *Streptococcus suis* (n = 14), *Streptococcus dysgalactiae* (n = 17), and *Streptococcus agalactiae* (n = 16). Glyco_hydro_25 was frequently observed in *Streptococcus pyogenes* (n = 9), *Streptococcus agalactiae* (n = 14), and *Streptococcus thermophilus* (n = 13) phage endolysins.

In terms of CWBD prevalence against bacterial hosts, ZoocinA_TRD was most frequent in *Streptococcus thermophilus* (n = 267), Choline_Bind_1 and Choline_Bind_3 in *Streptococcus pneumoniae* (n = 146, n = 85), SH3_5 in *Streptococcus pyogenes* (n = 47), *Streptococcus suis* (n = 22), and *Streptococcus dysgalactiae* (n = 25), SH3_3 in *Streptococcus agalactiae* (n = 13), CW_7 in *Streptococcus pyogenes* (n = 18) and *Streptococcus suis* (n = 28), and LysM in *Streptococcus suis* (n = 27) and *Streptococcus agalactiae* (n = 12). Figure 7 shows a heatmap of domain hit distributions across host species.

**Figure 7:**
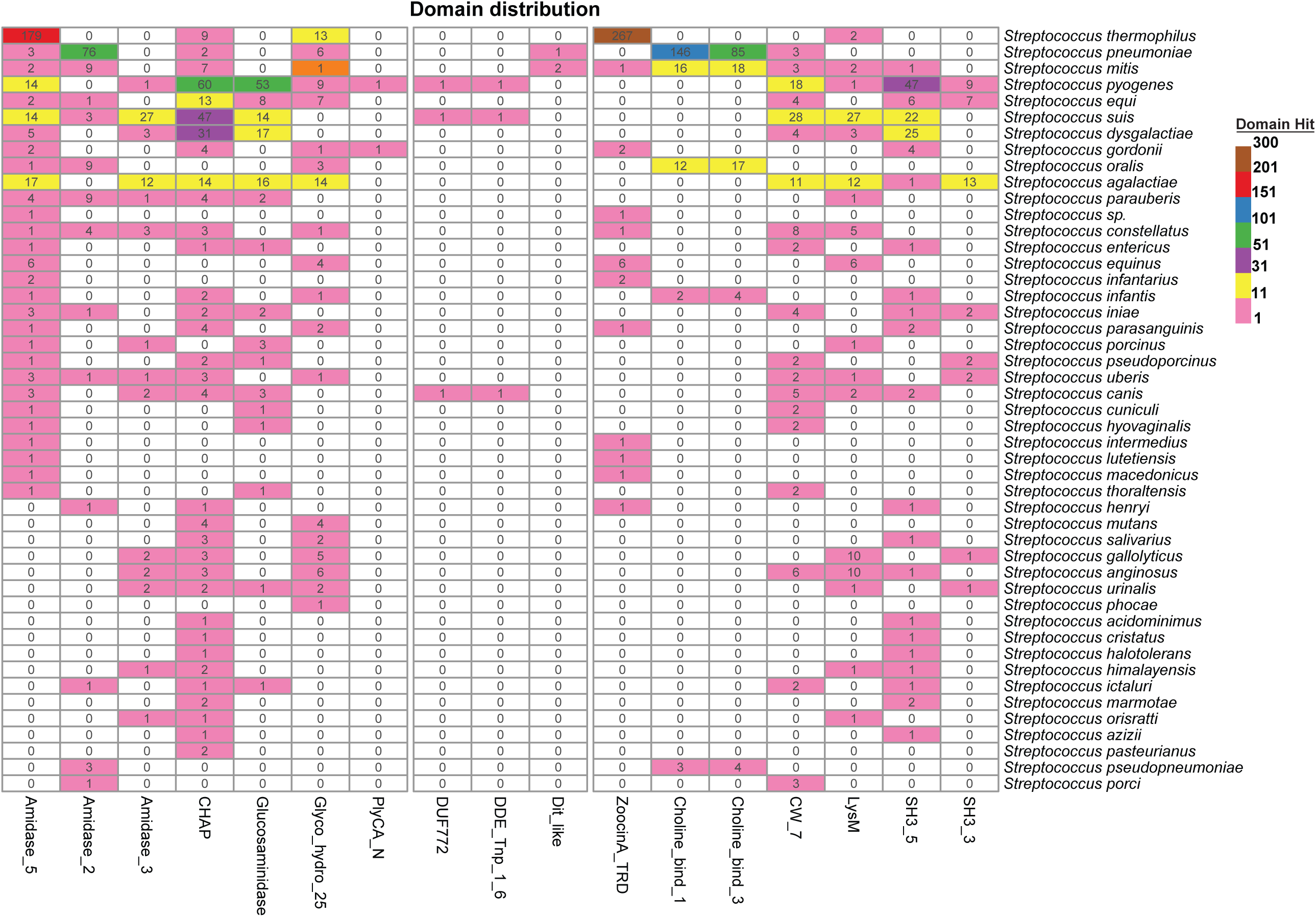
A heatmap illustrating the distribution of domain hit against bacterial hosts of *Streptococcus* phages. In X-axis, the EAD and CWBD are shown. Y-Axis indicates bacterial hosts. Cells are colored based on observed number of domain hits. The numbers within each cell indicate the number of hits for the corresponding bacterial host, whereas colors express a scale from lower (pink) to higher (Yellow brown) number of hits. Color range: Pink (1-10 hits), Yellow (11-30), Purple (31 – 50), Green (51 – 100), Blue (101-150), Red (151-200), yellow brown (201-300).

*Streptococcus* phages exhibit four types of endolysin gene organization: single gene, two adjacent genes, two spliced genes, and two distantly encoded genes (Figure 8). The majority (n=578) displayed single-gene organization. However, 63 phages, mainly *Streptococcus thermophilus* phages (n = 47), contained two adjacent genes: Amidase_5 and ZoocinA_TRD in one, ZoocinA_TRD alone in another. Similarly, 44 phages, primarily *Streptococcus thermophilus* phages (n = 40), showed two endolysin genes spliced by an endonuclease: Amidase_5 and ZoocinA_TRD, then ZoocinA_TRD. Nine phages, primarily *Streptococcus mitis* (n=4), *Streptococcus thermophilus* (n=2), and *Streptococcus dysgalactiae* (n=2), exhibited distantly encoded gene organization. Supplementary Table S10 summarizes the patterns of endolysin gene organizations. Phylogenetic analysis revealed that endolysins with the same domain architecture and pham do not typically form distinct monophyletic clades, with some exceptions (Figure 9).

**Figure 8:**
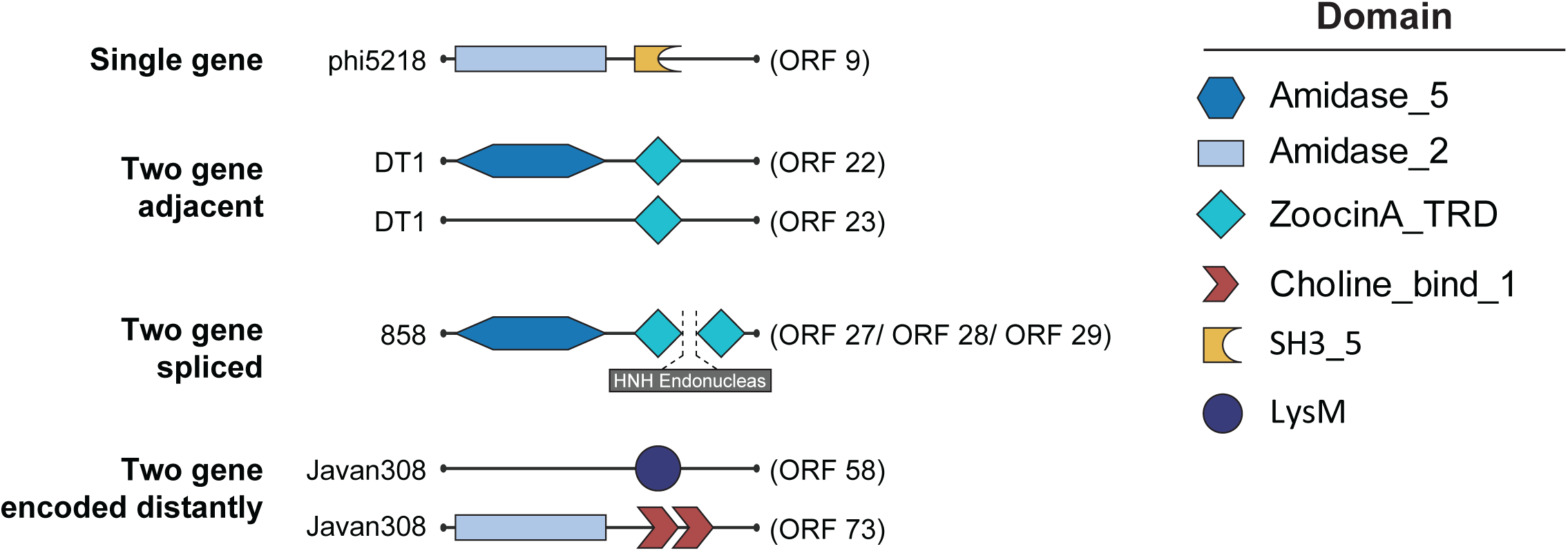
Observed endolysin gene organization strategy in *Streptococcus* phages. Four endolysin gene organization strategies were observed: single gene, two adjacent genes, two spliced genes, and two distantly encoded genes strategy. In phage phi5218, single gene (ORF 9) organization was observed and the endolysin showed Amidase_2 and SH3_5 domain architecture. In phage DT1, two gene adjacent organization was observed where ORF 22 had Amidase_5-ZoocinA_TRD domain architecture and ORF 23 only had ZoocinA_TRD domain. In phage 858 ORF 27 and ORF 29 were spliced by HNH endonuclease (ORF 28). In phage Javan308, two gene were encoded distantly where ORF 58 had only LysM domain and ORF 73 had Amidase_2 and two Choline_bind_1 domain.

**Figure 9:**
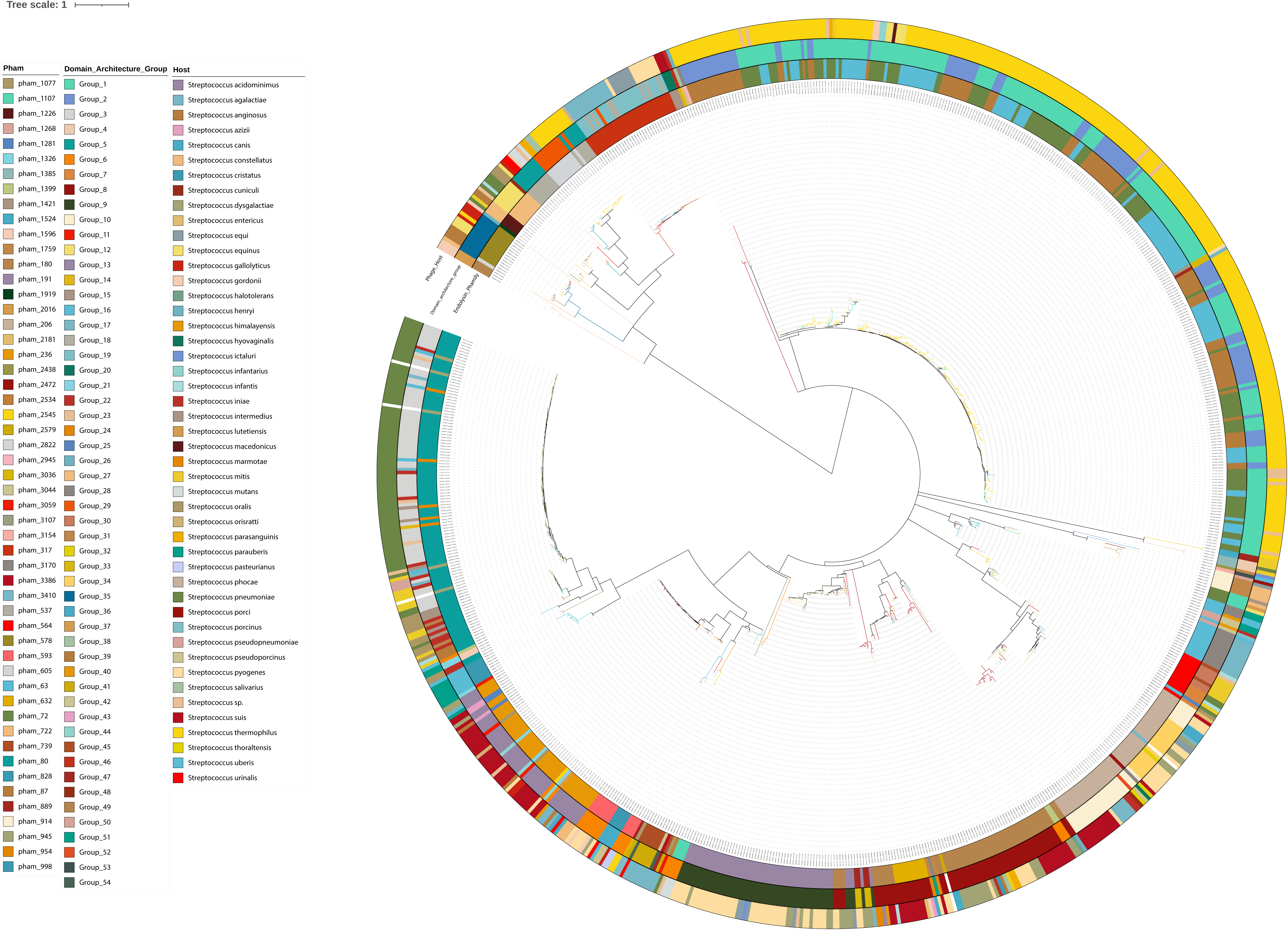
Phylogenetic tree of endolysin sequences of *Streptococcus* phages. The inner ring color strip represents endolysin phamily, middle ring represents Domain architecture-based group, the outer ring represents bacterial hosts of *Streptococcus* phages. Distinct monophyletic clade was not observed for endolysins belonging to same Domain architecture-based group or same endolysin phamily.

Endolysin phams with at least 10 unique protein sequences (63, 72, 80, 87, 180, 191, 206, 236, 317, 537, 564, 578) underwent adaptive/positive selection analysis using four tools (MEME, FUBAR, SLAC, FEL) from the Datamonkey web server. Each pham was screened for recombination events using GARD. Pham 578 (Glyco_hydro_25, LysM, LysM architecture) showed the highest recombination breakpoints, totaling 11. Positive/diversifying selection was identified at site 123 in pham 72 (Amidase_5, ZoocinA_TRD architecture), site 35 in pham 80 (Amidase_2, 2 Choline_Bind_1 repeat, 1 Choline_Bind_3 repeat), site 6 in pham 191 (Glucosaminidase, CHAP, SH3_5), and sites 11 and 202 in pham 206 (Amidase_5, CW_7, CW_7, Glucosaminidase) (Table S11).

### 3.4 Antimicrobial Peptide Identification and Characterizatio

814 endolysins were segmented into 20-mer peptides, yielding 268,194 fragments. After removing duplicates, 44,576 unique peptide fragments were analyzed for antimicrobial properties. Of these, 182 endolysin peptide (EP) fragments were consistently identified as antimicrobial by six prediction tools: AMPlify, amPEPpy 1.0, AxPEP, AmpGram, CAMPR4, and Antimicrobial Peptide Scanner vr.2. Results are summarized in Supplementary Table S12.

Among the 182 peptides, 16 were predicted to have antifungal properties by Antifp, while 66 were identified by iAMPpred (threshold cutoff = 0.8), with seven peptides exhibiting antifungal properties by both tools. Additionally, 20 peptides were predicted as antiviral by iAMPpred, whereas iACVP identified 135 peptides as antiviral, with 13 peptides recognized by both. To assess cell-penetrating properties, MLCPP 2.0 and BChemRF-CPPred were employed. MLCPP 2.0 predicted 32 peptides with cell-penetrating potential, 12 of which had high uptake efficiency; seven of these were from phage phiJH1301-3. BChemRF-CPPred identified 20 peptides with cell-penetrating properties, with eight peptides showing high uptake efficiency by both tools. ToxIBTL predicted most peptides (n = 167) as non-toxic. Supplementary Table S13 summarizes these findings.

### 3.5 Antimicrobial Peptide Stability and Molecular Docking

Alphafold2 was utilized to generate 3D structures for 182 antimicrobial peptides (AMPs), with Gromacs applied to assess their structural stability through molecular dynamics simulations. The average root-mean-square deviation (RMSD) for these structures ranged from 0.08 to 0.91 nm (Table S14). The peptide EP-60 (GVIESIFTHILGLCINAGLI) exhibited the smallest average RMSD of 0.08176, while EP-165 (GIKVVPYAPGIKPKAGWIFV) had the largest at 0.915646. Thirty AMP structures with low average RMSD values were selected for molecular docking with autolysin (PDB ID: 4X36). The binding site of autolysin identified by CASTp had an area of 173.225 Å² and a volume of 128.900 Å³. The HADDOCK score of these AMP-autolysin complexes varied between −79.6 ± 2.5 and −132.1 ± 2.2 (Table S15). Notably, EP-33 (IGETDYKKACHAIKAAGYAT), achieved the highest HADDOCK score of −132.1 ± 2.2, with an RMSD of 0.208 nm. The lowest HADDOCK score was for EP-152 (GLIESIFTHILGLCINTGLI), at −79.6 ± 2.5, with a lower RMSD of 0.135. Interestingly, EP-60 (GVIESIFTHILGLCINAGLI) had the lowest RMSD but a HADDOCK score of −89.7 ± 10.6. The top three scoring complexes—EP-33-autolysin, EP-121-autolysin (−128.0 ± 5.2), and EP-39-autolysin (−126.1 ± 3.2)—were selected for further molecular dynamics simulation analysis.

### 3.6 MD simulation of AMP-autolysin Complex

MD trajectories of AMP-autolysin complexes were analyzed for properties like RMSD, radius of gyration (Rg), and SASA (Figure 10). The EP-33-autolysin complex averaged an RMSD of 0.89 nm, stabilizing in the first 50 ns, peaking to 2.097 nm at 95 ns, and ending at 1.94 nm at 100 ns. The first 50 ns averaged 0.50 nm, rising to 1.29 nm in the last 50 ns. The EP-121 complex averaged 1.17 nm, peaking to 2.01 nm at 30 ns, with initial and final 50 ns averages of 1.06 and 1.28 nm, respectively, reaching 0.949 nm at 100 ns. The EP-39 complex had the lowest RMSD at 0.688 nm, peaking to 1.38 nm at 92 ns, with averages of 0.58 and 0.78 nm across each 50 ns interval, recording 1.15 nm at 100 ns (Figure 10A).

**Figure 10:**
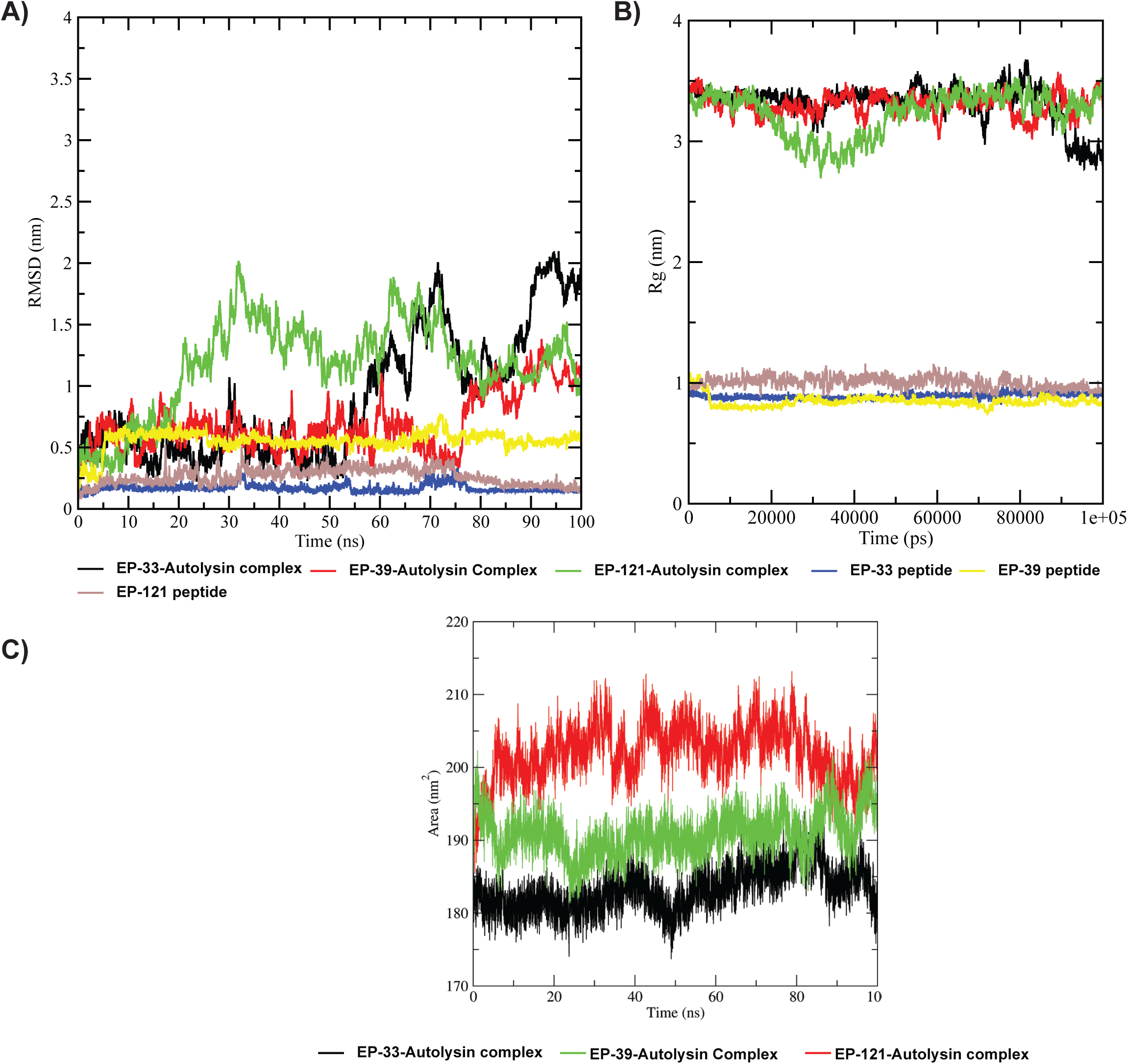
Trajectory analysis of protein-peptide complex over 100 ns molecular dynamics simulation. A) Root mean square deviation (RMSD) of Autolysin-Peptide complex and Peptides B) Radius of gyration (Rg) of Autolysin-Peptide complex and Peptides C) Solvent-accessible surface area (SASA) of Autolysin-Peptide complex.

The EP-33-autolysin complex showed an average Rg of 3.31 nm, peaking to 3.67 nm at 81.37 ns, ending at 2.77 nm. EP-121-autolysin complex averaged 3.23 nm, peaking to 3.54 nm at 82.1 ns, ending at 3.37 nm. EP-39-autolysin complex averaged 3.30 nm, peaking to 3.57 nm at 89.1 ns and recorded 3.30 nm at 100 ns (Figure 10B).

The EP-33-autolysin complex had the lowest average SASA value of 183.267 nm², peaking to 194.738 nm² at 85 ns, and measuring 181.267 nm² at 100 ns. The EP-121-autolysin complex had the highest average SASA value of 202.004 nm², peaking to 213.147 nm² at 78.83 ns, and measuring 201.105 nm² at 100 ns. The EP-39-autolysin complex had an average SASA value of 190.892 nm², peaking 202.286 nm² at 1 ns, with similar values at 85 ns and 95 ns (Figure 10C).

Using the MM/PBSA method, binding free energies over the last 30 ns revealed that the EP-121-autolysin complex had the most favorable binding energy of −39.31 Kcal/mol (Figure 11A), while EP-39 had the least at −20.71 Kcal/mol (Figure S9A), and EP-33 showed −35.26 Kcal/mol (Figure S9B).

**Figure 11:**
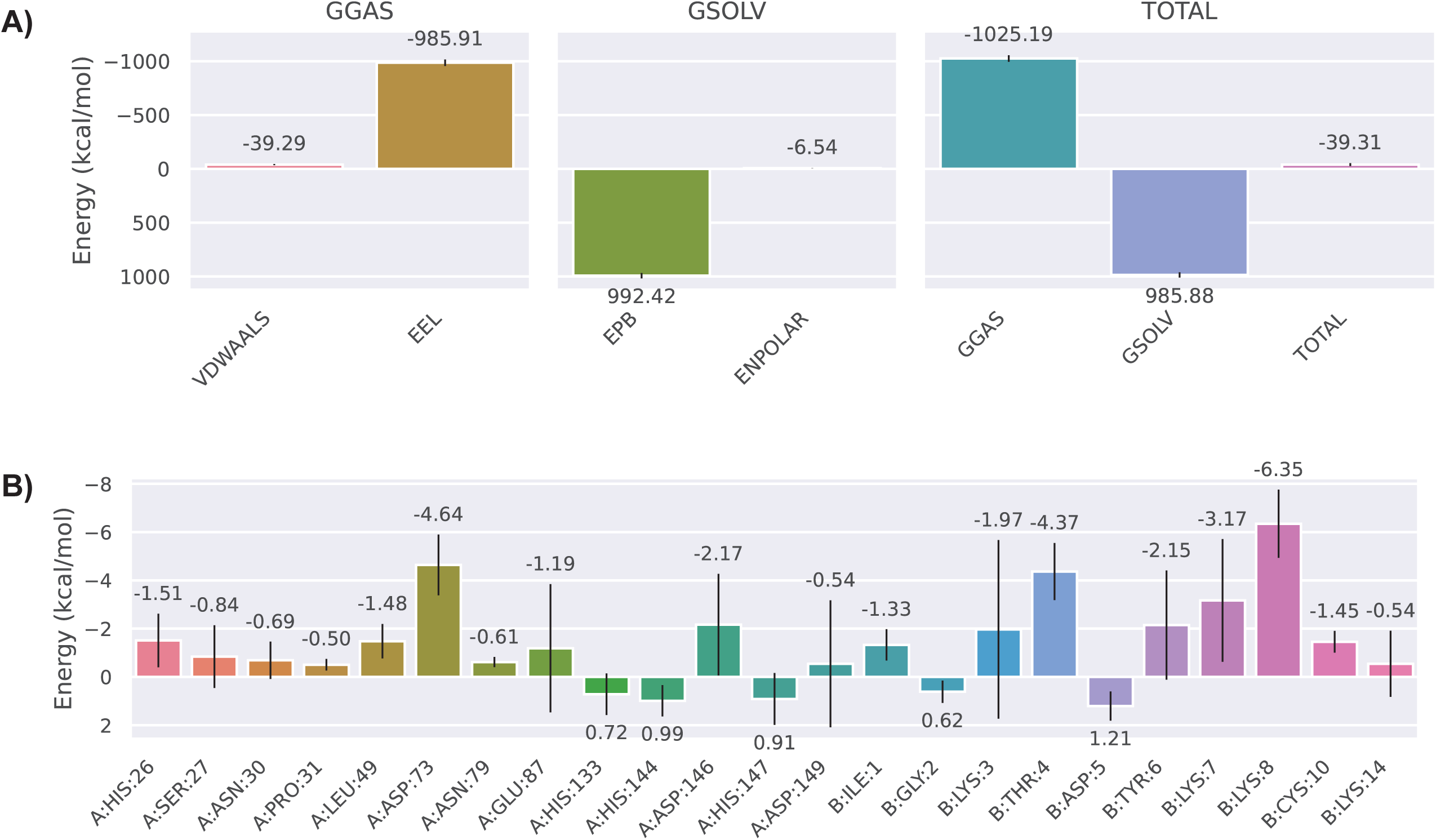
Binding free energy (Kcal/mol) analysis of EP-121-Autolysin complex by MM/PBSA method over last 30ns of 100 ns MD simulation. A) Energy components of binding free energy of EP-121-Autolysin complex. Here, VDWAALS and EEL indicate van der Waals interactions and electrostatic interactions in gas phase, respectively and GGAS indicates gas-phase energy. The solvation free energy (GSOLV) is also divided into two components, polar (EPB) and Non-polar (ENPOLAR). B) Per-residue binding free energy contribution of interacting residues of EP-121-Autolysin complex in Total Decomposition Contribution (TDC).

In the EP-121-autolysin complex, LYS8 of EP-121 had the highest contribution to the Total Decomposition Contribution (TDC) (−6.35 Kcal/mol), followed by ASP73 of autolysin (−4.64 Kcal/mol) and THR4 of EP-121 (−4.37 Kcal/mol). Residues like HIS133, HIS144, HIS147 of autolysin, and GLY2, ASP5 of EP-121 showed positive binding energy, indicating unfavorable binding (Figure 11B). A heatmap illustrating the change of per residue binding free energy of EP-121-autolysin complex is shown in Figure 12.

**Figure 12:**
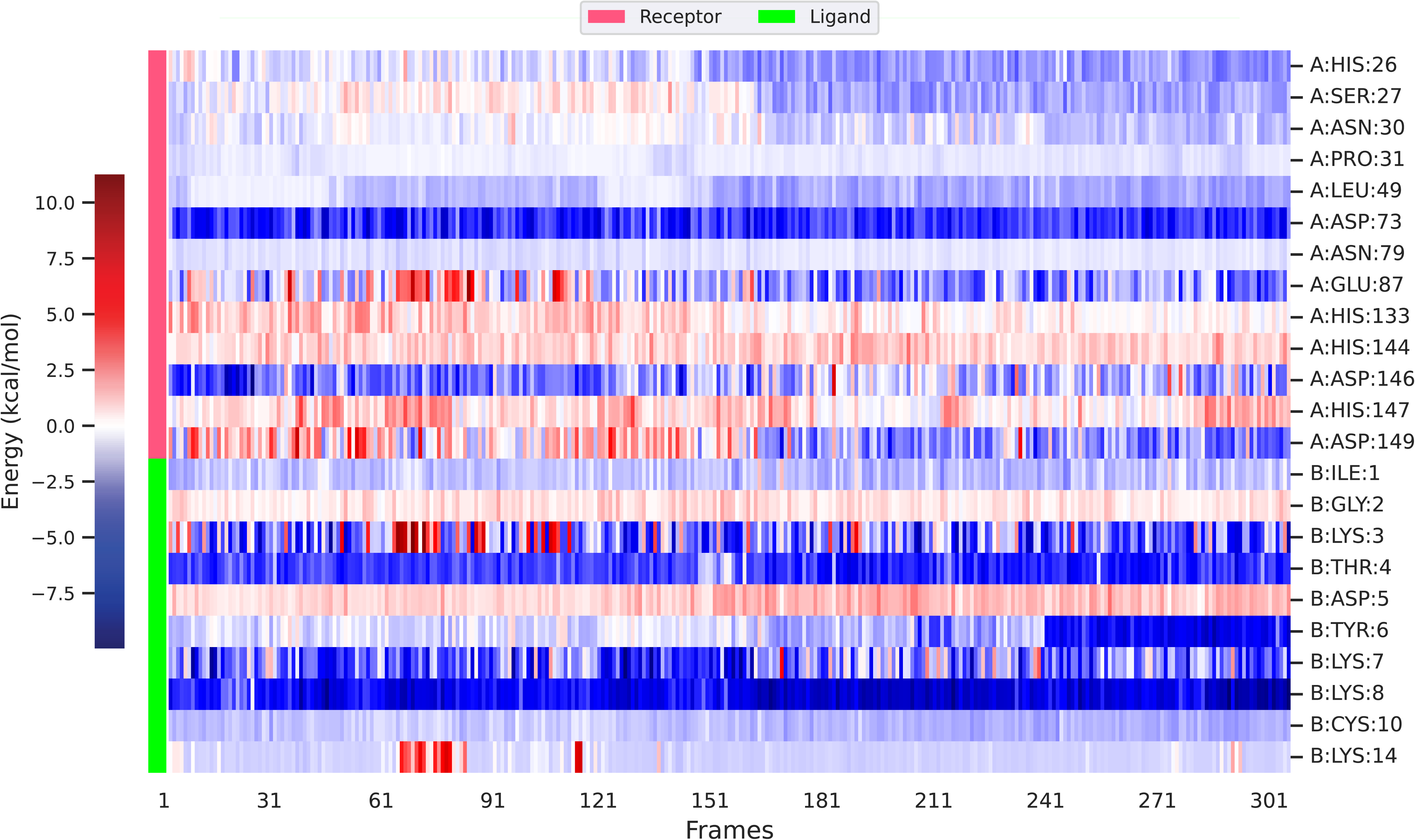
A heatmap illustrating the change of per-residue binding free energy contribution interacting residues of EP-121-Autolysin complex in Total Decomposition Contribution (TDC) over the last 30ns of 100 ns MD simulation.

In the EP-33-autolysin complex, TYR18 of EP-33 had the highest contribution of −6.62 Kcal/mol, followed by LYS14 (−6.45 Kcal/mol). Autolysin residues ASN30 (−1.97 Kcal/mol), HIS32 (−1.57 Kcal/mol), TRP72 (−1.58 Kcal/mol), VAL74 (−1.53 Kcal/mol), and HIS133 (−2.31 Kcal/mol) contributed favorably. In contrast, LYS45 (1.09 Kcal/mol), ASP73 (3.89 Kcal/mol), and ASP146 (1.47 Kcal/mol) indicated unfavorable binding, as did EP-33 residues LYS7 (2.22 Kcal/mol) and THR20 (1.56 Kcal/mol) (Figure S10A). In EP-39-autolysin complex, GLU87 of autolysin showed highest contribution (−4.07 Kcal/mol), followed by ARG19 (−3.85 Kcal/mol) and LYS16 (−3.22 Kcal/mol) of EP-39. Autolysin residues HIS26, LYS45, HIS133, and HIS147 showed positive binding energies (2.26, 1.85, 3.05, and 1.25 Kcal/mol), while EP-39 had a positive binding energy of 1.15 Kcal/mol at GLY20 (Figure S10B).

## 4. Discussion

Streptococcal diseases, caused by various *Streptococcus* species, range from mild infections to severe illnesses like pneumonia, bacteremia, and rheumatic fever. Rising antibiotic resistance in *Streptococcus* strains has intensified the search for alternative therapies, with endolysins and their antimicrobial peptides showing great potential (13,16,18,20,33,90,91). This study analyzes 709 *Streptococcus* phage genomes, revealing diverse endolysins and new taxonomic insights. Additionally, we explored antimicrobial peptides derived from *Streptococcus* phage endolysins and their interactions with the *S. pneumoniae* virulence factor autolysin.

Our results showed wide variability in *Streptococcus* phage genome lengths and GC content, ranging from 33% to 45%, consistent with host genomes. Interestingly, the presence of tRNA genes did not correlate with genome length, differing from other phages like those from *Pseudomonas* and *Vibrio cholerae* (37,39). For instance, phage Javan362, with a genome length of ∼41 kb, contains three tRNA genes. Among the 24 *Streptococcus* phages carrying ARGs, the majority were isolated from *Streptococcus suis* (n = 15) and *Streptococcus pyogenes* (n = 5), raising concerns about potential phage-mediated transduction of antibiotic resistance—a notable public health issue (92,93). Phages containing virulence genes were primarily isolated from *Streptococcus pneumoniae* (n = 73) and *Streptococcus pyogenes* (n = 66), possibly enhancing host fitness (94).

Clustering analysis of the phage genome identified 66 distinct *Streptococcus* phage clusters, with 35 singletons (4.93%), a lower rate compared to previous studies (34,37–39,95,96). We employed protein-sharing methods to cluster genomes, similar to approaches used in *Gordonia* and *Staphylococcus* phage studies (36,97), grouping proteins into phams using 35% amino acid identity and 80% coverage (35–38,40,97). Phylogenetic analysis supported these clusters, which generally formed distinct monophyletic clades, except for clusters 21, 41, and 64. Due to the lack of a universal core gene, we analyzed the core genes’ presence across clusters.

Our findings highlight inconsistencies in *Streptococcus* phage classification at both the family and genus levels. In 2022, ICTV classified 116 *S. thermophilus* phages into three genera— *Moineauvirus*, *Brussowvirus*, and *Vansinderenvirus*—within the family *Aliceevansviridae*.

However, proteomic tree analysis showed that these phages do not form a distinct monophyletic clade, a requirement for single-family assignment by ICTV (98). Specifically, 987 group phages (assigned to the *Piorkowskivirus* genus) formed a monophyletic clade with *Brussowvirus* phages and other phages from different *Streptococcal* hosts, sharing four core genes (99,100). *Moineauvirus* phages formed a monophyletic clade with phages from different clusters, sharing five core genes. *Vansinderenvirus* phages, previously known as 5093 group phages (101), formed a separate monophyletic clade, sharing seven core genes. The P738 group (unassigned) formed a distinct cluster with *Streptococcus pasteurianus* phages SG586P3 and SG586P1, sharing 38 core genes(102). Based on these findings, we propose revising the family assignment of phages within *Aliceevansviridae* and establishing a new family-level rank for *Streptococcus thermophilus* phages. We also suggest adding new members to the *Salasmaviridae* and *Rountreeviridae* families based on proteomic tree analysis and core gene presence. The proteomic tree and core gene sharing between inter-cluster members suggest the existance of numerous new families, following a similar approach recently applied to Lak Megaphages and Epsilon CrAss-like phages (103,104). Applying the ICTV’s 70% ANI criterion (98), we identified 296 genus subclusters and inconsistencies in the genus level assignment of *Moineauvirus*, *Brussowvirus* and *Piorkowskivirus* phages as they formed multiple genus clusters, while *Packlarkvirus* (subcluster 5A) and *Stonewallvirus* (subcluster 5D) formed single-genus cluster.

Our study uncovered significant diversity in the domain architectures of *Streptococcus* phage endolysins, reflecting their evolutionary complexity and therapeutic potential. This variation, particularly in domain arrangements, was linked to specific streptococcal hosts, suggesting adaptive evolution influenced by host specificity and environmental factors. Interestingly, despite similarities in domain architectures or sequence homology, phylogenetic analysis did not consistently reveal monophyletic clades for endolysins targeting particular streptococcal hosts. This suggests frequent domain exchange, gain, or loss among phages infecting different hosts, likely driven by homologous recombination during co-infection events or adaptive mutations in response to environmental pressures (105). The diversity observed across 47 *Streptococcus* hosts underscores the broad-spectrum potential of these endolysins against multidrug-resistant streptococcal strains. Our analysis revealed previously unreported domain architectures in *Streptococcus* phage endolysins (Figure 6), including CHAP domains, which had not been documented in pneumococcal endolysins until recently. Notably, the SP-CHAP endolysin, identified from an uncultured phage dataset, exhibited higher activity than Cpl-1 (106), while *S. pneumoniae* phages SP-QS1 and phage 33888 also possess CHAP domains, with the latter additionally harboring Choline_bind_1 and Choline_bind_3 domains. This discovery expands our understanding of endolysin diversity and offers new avenues for the development of chimeric endolysins like Cpl-7s, Cpl-711, and ClyJ (30–32).

A distinct domain combination of Amidase_5 and ZoocinA_TRD among *Streptococcus thermophilus* phages, often accompanied by a calcium-binding motif, suggested selective pressures from dairy environments favoring thermostable endolysins. This niche-specific selection likely contributed to reduced diversity in *S. thermophilus* endolysins compared to other streptococcal species (107). We also identified four gene organization patterns in *Streptococcus* phage endolysins—single-gene, adjacent two-gene, spliced two-gene, and distantly encoded two-gene, consistent with previous reports in *S. thermophilus* and *Staphylococcus* phages (36,108). Selective pressure analysis indicated that *Streptococcus* phage endolysins are under strong purifying selection, reflecting the evolutionary constraints that preserve their essential functions. However, we also identified specific sites under diversifying selection, which likely facilitate adaptation to new host environments (107).

In recent years, there has been a growing interest in antimicrobial peptides derived from endolysin, particularly for combatting resistant pathogens (91,109). Researchers have also been studying the potential synergistic effects of these antimicrobial peptides with endolysins and antibiotics (110–112). Our study identified 182 novel AMPs derived from *Streptococcus* phages, with some displaying antifungal, antiviral, cell-penetrating, and non-toxic properties. Molecular dynamics (MD) simulations were used to assess the stability of these peptides, and the top 30 stable peptides were subsequently docked with the *Streptococcus pneumoniae* virulence factor autolysin (*LytA*). Autolysin is crucial for *Streptococcus* pathogenicity, making it a potential target for new therapeutic approaches to combat streptococcal disease (113). Previous studies have shown that mutant pneumococcal isolates lacking autolysin exhibit minimal to no disease symptoms, underscoring its importance (114). Among the autolysin-AMP complexes, EP-39 displayed the lowest root-mean-square deviation (RMSD) value, indicating high complex stability, while the EP-121-autolysin complex had the most favorable binding energy according to end-state binding free energy analysis. Further experimental studies are needed to evaluate the therapeutic potential and safety of these peptides for treating bacterial infections in humans.

## 5. Conclusion

This study offers a comprehensive analysis of *Streptococcus* phages and their diverse lytic proteins (endolysins), uncovering their genomic diversity, providing novel taxonomic insights, and identifying new endolysin-derived antimicrobial peptides with the potential to target *S. pneumoniae*. Our study identified 66 distinct clusters of *Streptococcus* phages and 35 singletons, demonstrating variation in genome length, GC content, and shared gene content. Based on inter-cluster core gene sharing, average nucleotide identity (ANI) and a proteomic tree approach, we recommend a revision of *Streptococcus* phage taxonomy. This analysis proposes 21 new family-level classifications and 296 genus-level subclusters. The study also uncovered diverse domain architectures in *Streptococcus* phage endolysins, with some domains more prevalent in specific *Streptococcus* hosts, alongside previously unreported architectures. Additionally, the analysis revealed varying gene organization in endolysins, with purifying selection observed across most sites, except for some under-diversifying selection. This study is the first to report 182 novel *Streptocococcus* phage endolysin-derived antimicrobial peptides with potential activity against *Streptococcus pneumoniae*. These findings offer valuable insights into the genomic diversity of *Streptococcus* phages, highlighting the therapeutic potential of *Streptococcus* phage endolysins and their derived antimicrobial peptides as promising alternatives for combating *Streptococcus* infections.

## Acknowledgement

Not applicable.

## Ethical approval

Not required

## Supplemental materials

### Tables

**Table S1:** Information on the general characteristics of Streptococcus phages.

**Table S2:** Antimicrobial resistance genes identified by CARD in Streptococcus phage genomes.

**Table S3:** Virulence factors identified by VFDB in Streptococcus phage genomes.

**Table S4:** Proteomic Equivalence Quotient (PEQ) value between members within each cluster.

**Table S5:** Average nucleotide identity (ANI) value between members within each cluster.

**Table S6:** Detailed information related to each cluster.

**Table S7:** Information-related assignments of phages to proposed families and genus subclusters.

**Table S8:** key information on proposed family-level ranks, including shared core genes, genome lengths, GC content, coding capacity, and CDS counts.

**Table S9:** Information related to Endolysin grouping based on domain architecture.

**Table S10:** Information related to the patterns of endolysin gene organization, including adjacent genes, spliced genes, and distantly encoded genes.

**Table S11:** Information related to the selection pressure analysis on each endolysin pham (atleast 10 unique member).

**Table S12:** Information related to the results from all six AMP prediction tools for 182 endolysin-dervied antimicrobial peptides.

**Table S13:** Information related to Antifungal, Antiviral, toxicity and cell penetrating properties of 182 endolysin-derived antimicrobial peptides.

**Table S14:** Avearge RMSD value of 182 endolysin-dervied antimicrobial peptides obtained after 50ns MD simulation.

**Table S15:** HADDOCK result of top 30 AMP which showed lowest avearge RMSD value.

## Figures

**Figure S1:** Correlation between genome size of the phages (X-Axis) with GC percentage, Coding efficiency and Number of Coding sequences (CDS), each represented on the Y-axis. A) Correlation between genome size and GC % was negligible (R = 0.0029) B) the correlation between genome size and coding efficiency was negligible (R = 0.0033) C) In contrast, genome size vs CDS was moderately positive (R = 0.7286).

**Figure S2:** The phylogenetic tree generated by ViPTreeGen using 709 *Streptococcus* phage sequences along with all current members of *Caudoviricetes* except *Crassvirales* (n = 3,823, after duplicate removal, April 25, 2023). 21 new family level rank along with two previous family (*Salasmaviridae*, *Rountreeviridae*) could be suggested based on proteomic tree and inter-cluster shared core genes.

**Figure S3:** Genome comparison among members of cluster 18. The smallest *Streptococcus* phage Javan158 showed high conservation with Javan141.

**Figure S4:** Bar plot showing the number of orphams per cluster. Clusters (X-axis) are ordered in ascending order based on the number of orphams on the Y-axis.

**Figure S5:** Fragment of the phylogenetic tree generated by ViPTreeGen using 709 *Streptococcus* phage sequences along with all current members of *Caudoviricetes* except *Crassvirales* (n = 3,823, after duplicate removal, April 25, 2023) showing cluster 3 formed monophyletic clade with cluster cluster 42, 13, 14, and 31, sharing 4 core genes among 80 genomes.

**Figure S6:** Fragment of the phylogenetic tree generated by ViPTreeGen using 709 *Streptococcus* phage sequences along with all current members of *Caudoviricetes* except *Crassvirales* (n = 3,823, after duplicate removal, April 25, 2023) showing Cluster 12 formed a significant monophyletic clade with several other clusters, sharing 7 core genes among 94 members.

**Figure S7:** Bar plot depicting number of endolysins in each endolysin phamily. Endolysin phams (X-Axis) are ordered in ascending order based on the number of endolysins on the Y-axis.

**Figure S8:** Stacked bar plot showing distribution of observed domain architecture in endolysins against bacterial hosts of *Streptococcus* phages. X-axis indicates the number of occurrences of a specific domain architecture and Y-axis indicates bacterial hosts.

**Figure S9:** Binding free energy (Kcal/mol) analysis of EP-39-Autolysin complex (A) and EP-33-Autolysin complex by (B) MM/PBSA method over last 30ns of 100 ns MD simulation.

**Figure S10:** Per-residue binding free energy contribution of interacting residues of EP-39-Autolysin complex (A) and EP-33-Autolysin complex (B) in Total Decomposition Contribution (TDC).

